# Structure of the *Pseudomonas aeruginosa* PAO1 Type IV pilus

**DOI:** 10.1101/2024.04.09.588664

**Authors:** Hannah Ochner, Jan Böhning, Zhexin Wang, Abul K. Tarafder, Ido Caspy, Tanmay A. M. Bharat

## Abstract

Type IV pili (T4Ps), which are abundant in many bacterial and archaeal species, have been shown to play important roles in both surface sensing and twitching motility, with implications for adhesion, biofilm formation and pathogenicity. While Type IV pilus (T4P) structures from other organisms have been previously solved, a high-resolution structure of the native, fully assembled T4P of *Pseudomonas aeruginosa,* one of the major human pathogens, is not available. Here, we report a 3.2 Å-resolution structure of the *P. aeruginosa* PAO1 T4P determined by electron cryomicroscopy (cryo-EM). PilA subunits constituting the T4P exhibit a classical pilin fold featuring an extended N-terminal α-helix linked to a C-terminal globular β-sheet-containing domain, which are packed tightly along the pilus. The N-terminal helices constitute the pilus core where they stabilise the tubular assembly via hydrophobic interactions. The α-helical core of the pilus is surrounded by the C-terminal globular domain of PilA that coats the outer surface of the pilus, mediating interactions with the surrounding environment. Comparison of the *P. aeruginosa* T4P with T4P structures from other organisms, both at the level of the pilin subunits and the fully assembled pili, allows us to enumerate key differences, and detect common architectural principles in this abundant class of prokaryotic filaments. This study provides a structural framework for understanding the molecular and cell biology of these important cellular appendages mediating interaction of prokaryotes to surfaces.

## Introduction

Surface sensing and adhesion are crucial for bacteria and archaea to colonise new environments^1,2^ as they help cells invade hosts, establish biofilms and form connections in microbiomes^3–6^. These sensing and adhesion processes are often mediated by proteinaceous, filamentous surface structures known as pili or fimbriae^7–10^. Bacterial surface pili are assembled by repeated interactions of monomeric protein subunits called pilins, leading to the formation of a filamentous structure that protrudes from the cell surface and binds to substrates^11,12^.

Although there is a bewildering variety of pili present on bacteria, the Type IV pili (T4Ps) of Gram-negative bacteria are profoundly involved in surface sensing and initial attachment of bacteria to surfaces^13^. Initial attachment to surfaces is not only the first step of biofilm formation^14^, but is also important in pathogenicity and the establishment of infections^15^. T4Ps are dynamic filaments that can extend and retract^16,17^, allowing cells to rapidly respond to a changing environment, particularly when encountering a favourable surface suitable for colonisation.

*Pseudomonas aeruginosa* is an important Gram-negative human pathogen that poses a critical problem in hospital settings^18^, causing widespread antibiotic-resistant infections^19^. *P. aeruginosa* evades antibiotic treatments by forming biofilms^20,21^, which makes clearing bacterial infections extremely challenging. Biofilm formation in *P. aeruginosa* is facilitated by the expression of a variety of filamentous adhesins^6,22^ and pili^13,14^ on its outer surface. As *P. aeruginosa* T4Ps mediate surface sensing^14^, the first step of biofilm formation, they play a key role in allowing *P. aeruginosa* to transition from a unicellular planktonic to a multicellular biofilm state. Understanding the structure of the *P. aeruginosa* Type IV pilus (T4P) is thus highly relevant for the development of strategies to combat antibiotic-resistant infections.

On the structural level, several studies have reported the structures of the filamentous T4Ps from a variety of prokaryotes, including *Neisseria gonorrhoeae, Escherichia coli, Myxococcus xanthus, Pyrobaculum arsenaticum* and *Thermos thermophilus*^23–29^ using electron cryomicroscopy (cryo-EM) single-particle analysis. Other studies have elucidated the organization of the secretion machinery of T4Ps, called Type IV secretion systems (T4SS), in the periplasm of different bacterial species using electron cryotomography (cryoET)^23,30–32^. However, the high-resolution structure of the filamentous T4P from the model pathogen *P. aeruginosa* has remained elusive. While previous studies have reported an X-ray crystal structure of the pilin monomer from *P. aeruginosa* strain K (PAK)^33^, the arrangement of the full pilus from the monomer subunits could not be inferred. Additionally, a low-resolution cryo-EM structure of the *P. aeruginosa* PAK pilus was reported, which could not be improved due to structural heterogeneity confounding image processing^24^.

In this study, we describe a 3.2 Å-resolution cryo-EM structure of the fully assembled *P. aeruginosa* T4P from the model PAO1 strain. Our structure allows us to derive an atomic model for the pilus, which provides detailed molecular insights into the assembly and three-dimensional arrangement of this important appendage. Furthermore, our structural data allows a detailed comparison of this pilus with other solved T4P structures from multiple bacterial species, which is relevant for placing past and future structural and cell biology studies of this abundant pilus into context.

## Results

### Purification of natively assembled T4Ps

To study the structure of the *P. aeruginosa* T4P, we used a PAO1 strain with a deletion of the retraction ATPase *pilT*^13,17,34,35^ to maximise pilus decoration on the cell surface. Cells from this strain were deposited on electron microscopy grids and visualised with cryoET (Methods, Fig. 1a). Multiple T4Ps were observed protruding from the outer membrane of these PAO1 Δ*pilT* cells, with the same overall morphological appearance as T4Ps described in other bacteria^36^.

**Figure 1:**
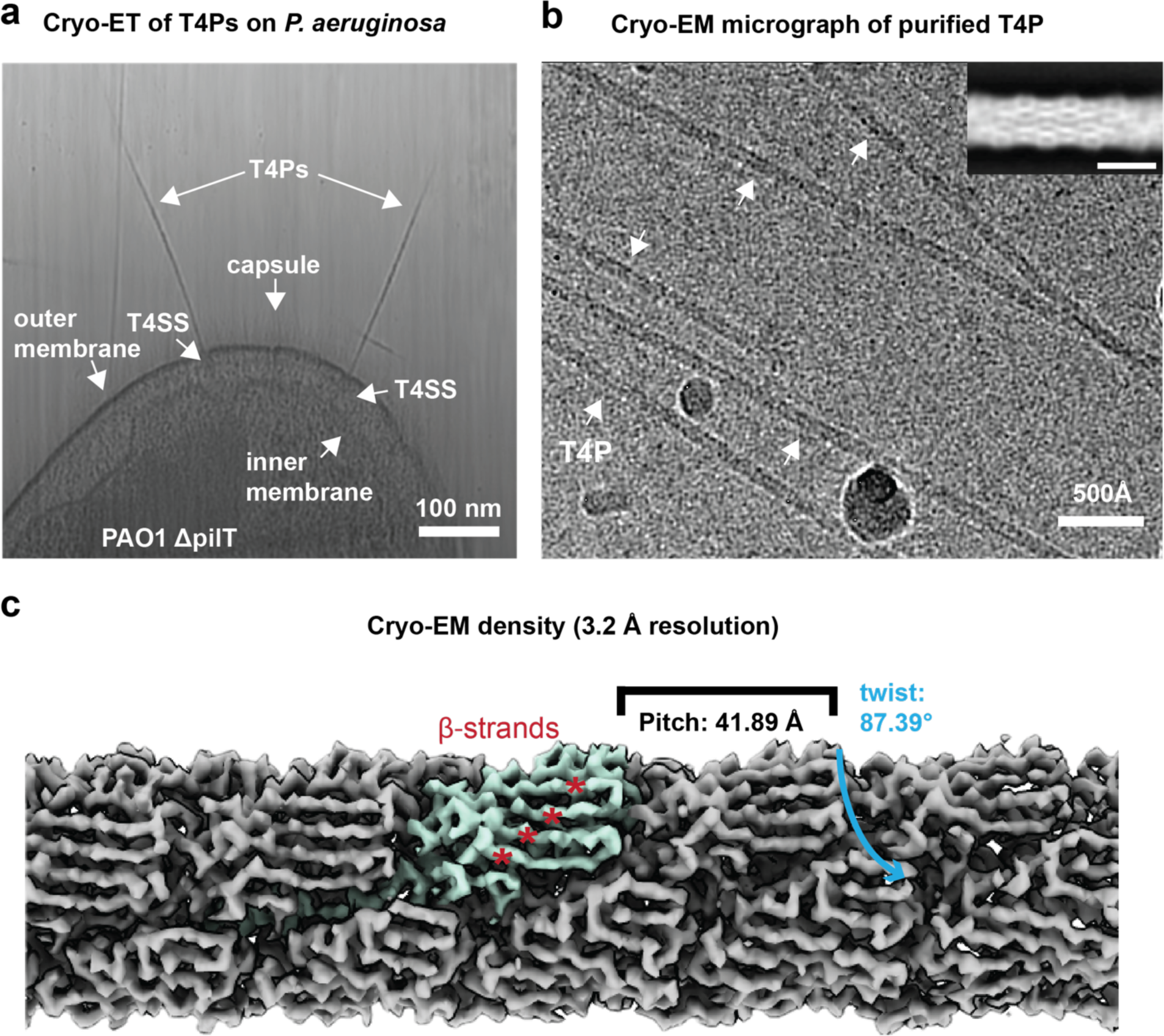
Cellular arrangement and structure of the *P. aeruginosa* PAO1 T4P. **a** Slice through a tomogram of a *P. aeruginosa* PAO1 τι*pilT* cell with T4Ps protruding from the cell. Cell membrane, capsule and T4Ps, as well as T4SS, which assemble the T4Ps, are marked by arrows. **b** Cryo-EM micrograph of T4Ps purified from *P. aeruginosa* PAO1 τι*pilT* cells, with T4Ps labelled by white arrows. The inset shows a representative 2D class average (scale bar 50 Å). **c** Cryo-EM density map of the assembled T4P reconstructed in RELION 4.0 with one of the PilA subunits highlighted in turquoise. The map resolution of 3.2 Å shows a clear separation of the β-strands in each of the PilA subunits (marked by red asterisks). TheT4P has a helical rise of 10.17 Å, a helical twist of 87.39° and thus a helical pitch of 41.89 Å.

The increased occurrence of T4Ps on the cell surface of the PAO1 Δ*pilT* strain facilitated pilus isolation for structural analysis. We adapted a previously described methodology for purifying T4Ps^37^ to isolate *P. aeruginosa* T4Ps in this study. In our preparations, along with the T4Ps sheared from cells, bacterial flagella were present as contaminants (Fig. S1). For cryo-EM analysis, the contaminant flagella did not pose an unsurmountable problem because of the large difference in diameter of the two filaments (flagella: ∼170 Å^38^, T4P: 51 Å), which allowed us to clearly distinguish them in micrographs. Peptide fingerprinting mass spectrometry confirmed the presence of *P. aeruginosa* PilA, the protein constituting the T4P subunits, in our preparation with 70% coverage of the PilA sequence (Fig. S1).

### Structure of the *P. aeruginosa* PAO1 T4P using cryo-EM helical reconstruction

Once the presence of natively assembled T4Ps was confirmed in our sample, we proceeded to acquire cryo-EM data on this specimen (Fig. 1b), which allowed us to produce two-dimensional (2D) class averages of the *P. aeruginosa* PAO1 T4P (Fig. 1b, inset). The 2D class averages showed a tubular assembly with a diameter of approximately 51 Å.

Next, we used a previous estimate of the helical symmetry of the *P. aeruginosa* PAK strain T4P^24^, whose PilA subunit has 67% sequence identity with the PAO1 PilA, as a starting point, and performed repeated three-dimensional classification and refinement (Fig. S2). Particles from classes not showing densities that resembled protein subunits were discarded. The remaining classes, which exhibited significantly improved image processing statistics and cryo-EM densities, were retained for further processing. The helical symmetry of these classes was applied to the remaining dataset in subsequent refinements, which allowed us to perform helical reconstruction and obtain a 3.2 Å-resolution cryo-EM map (Fig. 1c), from which an atomic model of the T4P could be built (Table S1, Movie S1, and Figs. 2 and S3-4).

**Figure 2:**
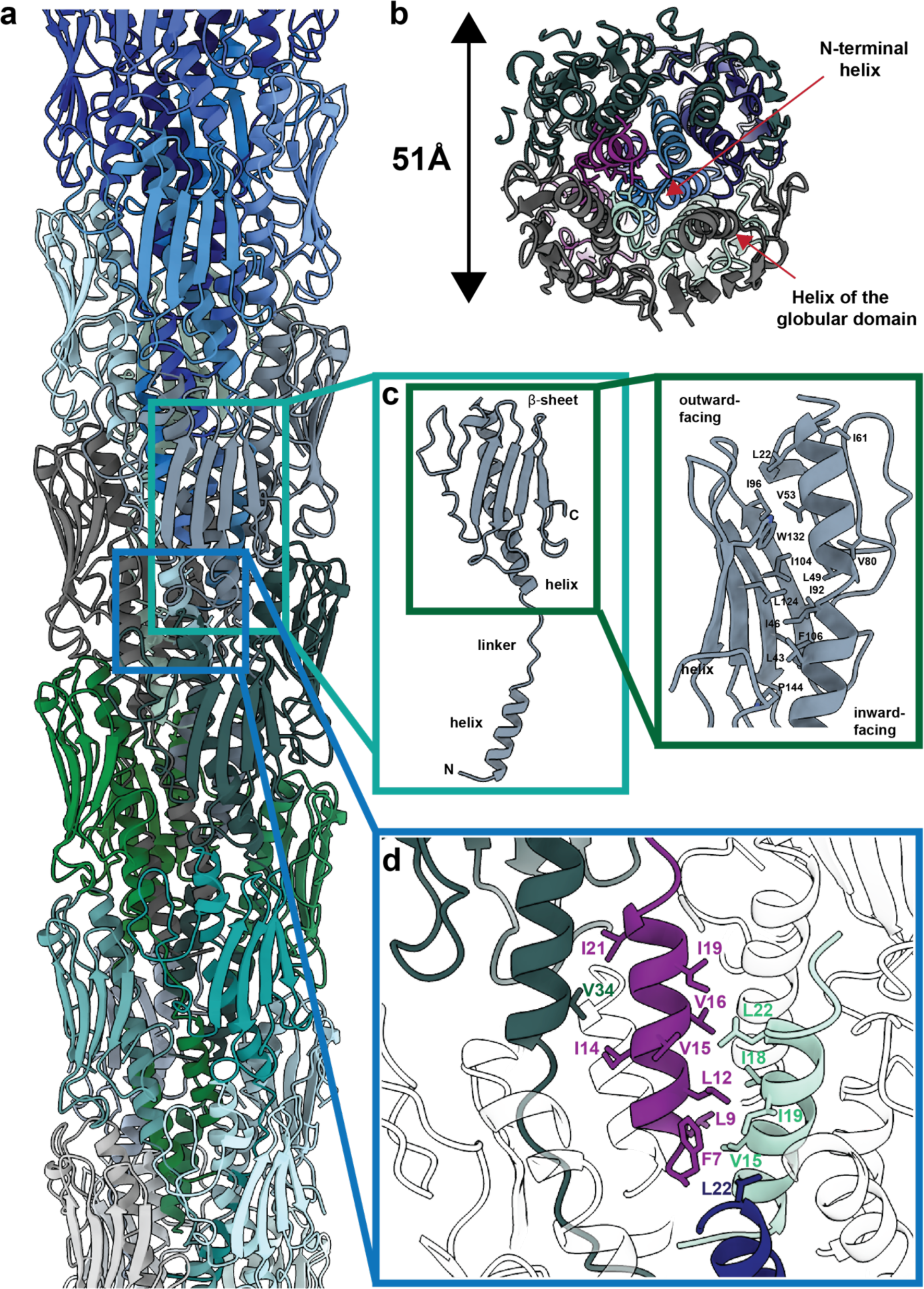
Structural features of the *P.aeruginosa* PAO1 T4P. **a** Model of the assembled *P. aeruginosa* PAO1 T4P. **b** Cross section through the model showing the tightly packed N-terminal α-helices in the centre of the pilus, which in turn interact with a helix in the globular domain of another pilin subunit (marked by red arrows). **c** Structure of an individual PilA subunit composing the pilus. The PilA subunit features an N-terminal α-helix and a globular domain consisting of an α-helical segment connected to a β-sheet by a linker. The two helices (N-terminal and globular domain α-helices) are coupled by a ‘melted’ linker region. N- and C-termini are marked N and C, respectively. The inset highlights the strong hydrophobic interactions between the α-helix and β-sheet in the globular domain, which stabilise the pilin structure. **d** Hydrophobic subunit-subunit interactions in the core of the T4P are mediated by the PilA α-helices.

The atomic model confirms a tubular arrangement of the *P. aeruginosa* PAO1 T4P (Fig. 2a), with a helical rise of 10.17 Å and a right-handed twist of 87.4° per subunit, resulting in an overall helical pitch of 41.9 Å (Fig. 1c and 2a). In agreement with classical T4Ps, the N-terminal α-helix of each PilA subunit is stacked in the centre of the tubular T4P to form the filament (Figure 2a-b). This N-terminal helix is connected through a short linker to C-terminal globular pilin domain (Fig. 2c). The globular C-terminal pilin domain contains a β-sheet consisting of four β-strands that face the exterior of the T4P and are thus exposed to the solvent (Fig. 2c). These β-strands are stabilised by a long α-helix containing several hydrophobic residues that interact with β-sheet residues along the T4P tube (Fig. 2c and S5).

The innermost part of the T4P is stabilised by hydrophobic stacking of the N-terminal α-helices of PilA, containing a row of hydrophobic residues (amino acid residues 7-22, where residue 7 is the N-terminus of the mature pilin), forming the core of the T4P (Fig. 2d). These N-terminal helices in turn interact with the long helices of the globular domain through further hydrophobic interactions (Fig. 2b, d). Overall, these interactions result in an ordered, tubular filament (Fig. 2a).

Previous work has shown that five groups of T4Ps are present in *P. aeruginosa* strains, with each strain expressing a single group. Two of these groups are glycosylated (Group 1 and Group 4), which are modifications that help bacteria escape phages that use the T4P as an entry receptor^39–41^. *P. aeruginosa* PAO1, however, is a strain that encodes a Group 2 pilin that is not glycosylated, in good agreement with our cryo-EM density which did not show any densities that are not accounted for by the polypeptide chain itself (Movie S1).

### Comparison of T4P structures

The PilA subunit of the *P. aeruginosa* PAO1 T4P adopts a classical pilin fold, as demonstrated by comparison with pilin subunits from other organisms (Figure 3a-g). While Type IV pilins generally consist of a globular domain and an N-terminal helix, the globular domain is considerably less conserved than the N-terminal helix (Figure 3h), which is a major mediator of subunit-subunit interactions (Figure 2d), stabilizing the core of the pilus. In addition, the C-terminal loop contains two buried cysteine residues (C134, C147) in close proximity, which form a disulfide bond resolved in our map, constituting a structural feature present in many known T4P structures^42^.

**Figure 3:**
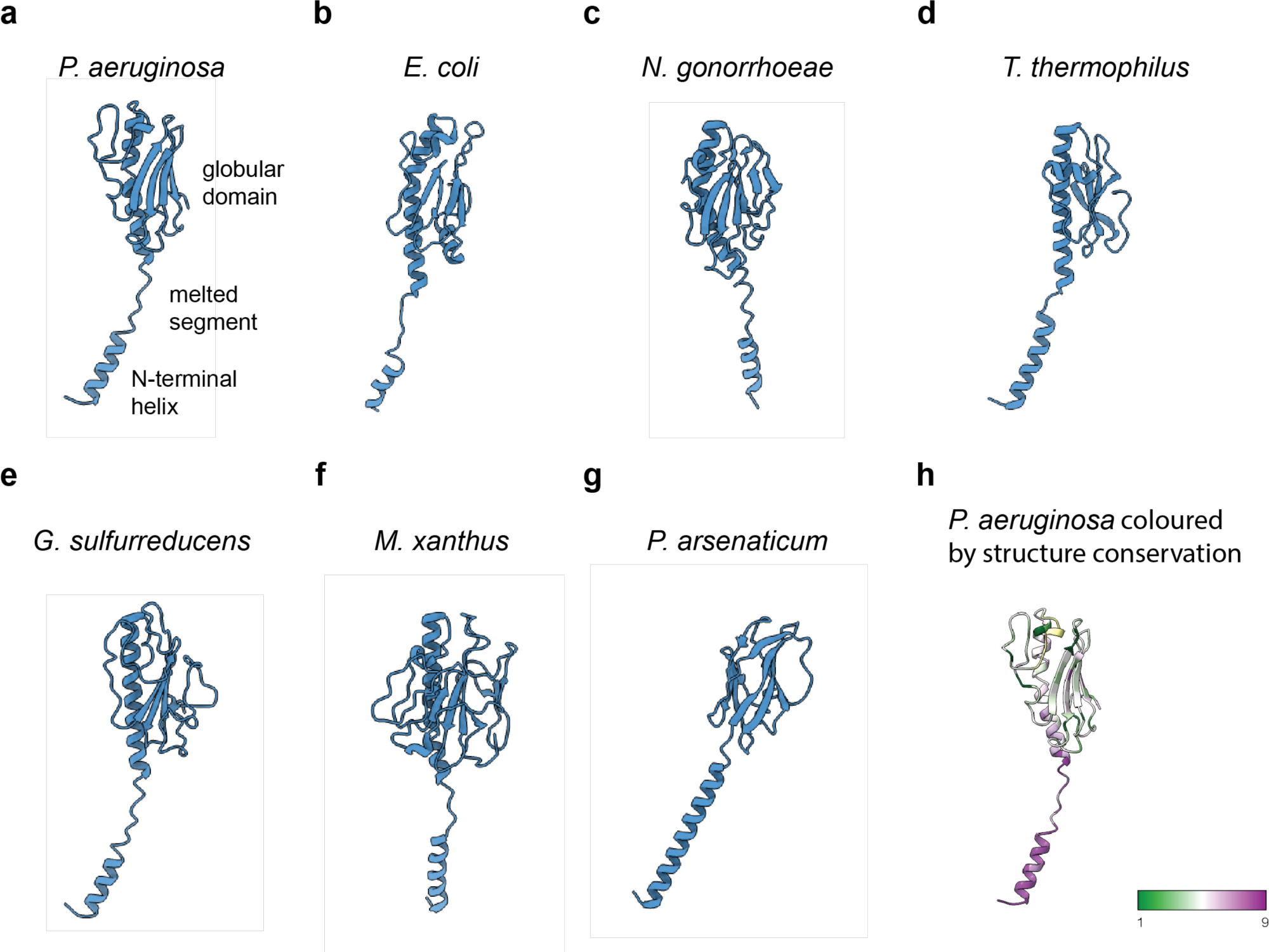
Comparison of Type IV pilin structures from different organisms. **a-g** Aligned structures of Type IV pilins from different organisms: *P. aeruginosa* PAO1 (this study, PDB: 9EWX) (**a**), *E. coli* (PDB: 6GV9) (**b**), *N. gonorrhoeae* (PDB: 5VXX) (**c**), *T. thermophilus* (PDB: 6XXD) (**d**), *G. sulfurreducens* (PDB: 6VK9) (**e**), *M. xanthus* (PDB: 8TJ2) (**f**), *P. arsenaticum* (PDB: 6W8U) (**g**). The pilin structures have been aligned to the *P. aeruginosa* pilin using the ChimeraX *matchmaker* function. The resulting orientations show the structural diversity between the pilins introduced by the presence of the melted linker region featured by most pilin structures. **h** *P. aeruginosa* PilA pilin ribbon diagram coloured based on structure conservation score from highly conserved (9) to low conservation (1) calculated by comparing the pilin structures presented in **a-g** by sequence alignment using ConSurf^74–76^. While the N-terminal α-helix is highly conserved, the β-sheet part of the pilin exhibits low conservation.

While early X-ray structures of PilA from other *P. aeruginosa* strains (such as the PAK strain) suggested that the N-terminus of the PilA subunit forms a continuous helix spanning the full length of the subunit^43,44^ (Fig. S6), in subsequent cryo-EM structures – and in our 3.2 Å-resolution structure - a non-helical but well-resolved linker segment (Fig. 2c, S4b) containing a prominent helix-breaking proline residue (P28, Fig. S4b), which divides the α-helical part of PilA into an N-terminal helix (residues 7-22) and a distinct α-helix that is part of the globular domain (residues 31-58). This so-called ‘melting’ of the middle of the N-terminal helix is important for assembly of the pilus^24^ and is broadly conserved among bacterial pilins (Fig. 3a-f), while more divergent organisms, such as the archaeon *Pyrobaculum arsenaticum* (Fig. 3g), exhibit a more extensive N-terminal helix without melted segments.

Next, we performed an overall structural comparison between described Type IV pilins from various prokaryotic organisms. While the globular domains are similar at an organizational level, the presence of the ‘melted’ linker segment results in a different positioning of the N-terminal helix relative to the globular domain, influencing overall pilus architecture (Table S2). This becomes especially clear when aligning the pilin structures as shown in Figure 3a-g, where each of the pilin models is presented in an orientation resulting from structural alignment to the *P. aeruginosa* pilin structure (Fig. 3a) using the *matchmaker* function in ChimeraX^45^.

A comparison of T4P structures originating from different organisms is not only of interest on the level of the pilin structure, but also when considering the overall architecture of the fully assembled pili (Figure 4). All T4Ps present an overall straight, tubular geometry; most also feature similar helical parameters (rise 9-11 Å, twist 87-102°, see Table S2), the only notable exception being the archaeal pilus structure of *P. arsenaticum*, which has a much smaller helical rise (5.3 Å) and pitch (compare 19 Å in *P. arsenaticum* with 36-43 Å in other pili). Despite the relatively similar helical symmetry amongst bacterial T4Ps, the pilus structures differ significantly in diameter, ranging from 51-80 Å. The *P. aeruginosa* T4P shows the smallest diameter (51 Å) by a large margin (Table S2). This comparatively small diameter is remarkable when combined with the observation that the *P. aeruginosa* T4P does not present flexible loops on the outer surface of the pilus, which are observed prominently in the structures of other pili (Fig. 4, especially 4c-e, g, marked by red arrows). As flexible residues could potentially be targets for extracellular proteases, a lack of such residues might make the *P. aeruginosa* T4P more resistant to proteolysis. This is likely important in the harsh and diverse environments that *P. aeruginosa* must encounter during biofilm formation and infection.

**Figure 4:**
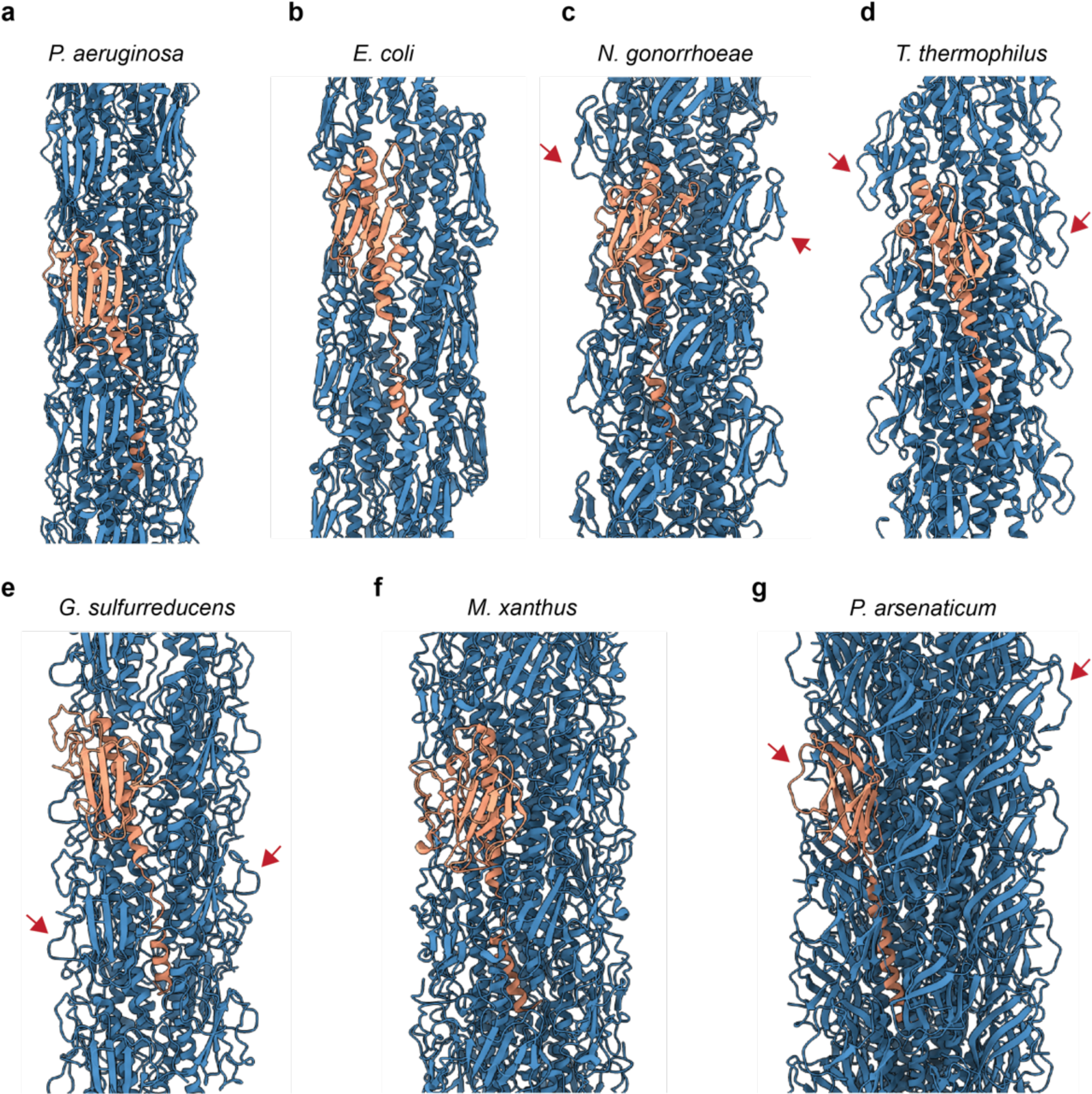
Comparison of T4P architectures in different organisms. **a-g** Models of T4P structures from different organisms with one pilin subunit highlighted in orange for each structure: *P. aeruginosa* PAO1 (this study, PDB: 9EWX) (**a**), *E. coli* (PDB: 6GV9) (**b**), *N. gonorrhoeae* (PDB: 5VXX) (**c**), *T. thermophilus* (PDB: 6XXD) (**d**), *G. sulfurreducens* (PDB: 6VK9) (**e**), *M. xanthus* (PDB: 8TJ2) (**f**), *P. arsenaticum* (PDB: 6W8U) (**g**). Flexible loops on the pilus surface are marked by red arrows.

## Discussion

In this study, we present the structure of the T4P from the important human pathogen *P. aeruginosa.* Past studies on the *P. aeruginosa* T4P from a different strain (PAK strain), had reported a low-resolution cryo-EM map, from which an atomic model could not be directly derived^24^. With improved methods for cryo-EM^46^, image analysis^46–52^ and helical three-dimensional classification^53–57^, we were able to deduce the helical symmetry of the *P. aeruginosa* T4P PAO1 strain and resolve its structure (Figs. 1-2).

Our structure reveals that the *P. aeruginosa* PilA protein adopts a classical pilin fold, sharing similarities with other pilins across the prokaryotic tree of life (Fig. 3). The N-terminal α-helix of PilA extends away from the body of the pilin globular domain to stack with other pilin subunits, mediating major subunit-subunit interactions within the pilus. Such a ‘hydrophobic arm’-like element is also commonly found in other pili where it similarly provides extensive contact with hydrophobic regions of other subunits, for example in filaments employing donor-strand exchange^53,58,59^. A unique feature of T4Ps is the simultaneous interaction of one N-terminal region with various regions of several other subunits within the pilus (Figs. 2). As a result of the interactions discussed above, PilA monomers of *P. aeruginosa* PAO1 stack in three-dimensional space to form a tubular T4P, resembling pili from other species (Fig. 4). While the overall architecture of the T4Ps is similar, they differ significantly in diameter and in the details of their outer surface structure.

The base of the T4P filament solved in this study interacts with T4SS in the periplasmic space of *P. aeruginosa* cells. Although the T4SS of other bacteria have been studied structurally^30,31^, characterization of the T4SS of *P. aeruginosa* at the structural level remains an open avenue of research. The other end of the pilus contains an important *P. aeruginosa* adhesin called PilY1^60^. While PilY1 should be retained during purification, the cryo-EM data of pili tips exhibited a strong structural heterogeneity, which hindered convergence to a convincing density for the PilY1-bound tip. Therefore, the question of how PilY1 interacts with and decorates the pilus tip remains unresolved. Further studies on native cellular systems, i.e. cells and biofilms, will be required to determine the exact mode of interaction between different T4P components^61^.

Our structure will serve as a reference for future molecular and cellular studies on this T4P from a major human pathogen. However, further research needs to be performed on the *P. aeruginosa* T4P to understand its precise mechanistic role in mediating surface adhesion, motility and human infection.

## Materials and Methods

### Isolation of T4Ps from *P. aeruginosa*

T4Ps were isolated and purified following a protocol adapted from Rivera *et al*^37^. Briefly, *P. aeruginosa* PAO1 Δ*pilT* (kind gift from Urs Jenal, University of Basel) was streaked from a glycerol stock on Lysogeny broth (LB)-agar plates and incubated overnight at 37 °C. Four single colonies from these plates were streaked onto new LB-agar plates and grown overnight at 37 °C. Cells were scraped from these plates and resuspended in LB media to obtain a suspension with an optical density (OD_600_) of 8.0. Fifty μl of this suspension was spread onto 30 LB-agar plates, which were incubated overnight at 37 °C until bacterial lawns were formed. Lawns were harvested in 5 ml ice-cold buffer A per plate (150 mM ethanolamine pH 10.5, 1 mM dithiothreitol) and incubated for 1 hour at 4 °C with stirring. To shear T4Ps from cells, samples were transferred to 50 ml Oakridge centrifuge tubes (Beckman Coulter) and vortexed three times at maximum strength for 1 minute at a time with 2-minute intervals on ice in between. To remove cells, samples were centrifuged at 15,000 relative centrifugal force (rcf) for 30 minutes at 4 °C, supernatants collected and centrifuged again at 15,000 rcf for 10 minutes at 4 °C. The supernatants from the second centrifugation step were then dialysed overnight at 4 °C against ice-cold buffer B (50 mM Tris pH 7.5, 150 mM NaCl) in SnakeSkin dialysis tubing (Molecular weight cutoff 3 kDa, Thermo Fisher Scientific) until the sample reached pH 7.5, which was confirmed using pH strips. To harvest pili, the sample was centrifuged at 20,000 rcf for 40 minutes at 4 °C. The resulting pellet containing T4Ps was resuspended in buffer B. T4Ps were further purified by adjusting the solution to 0.5 M NaCl, followed by T4P precipitation by addition of 10% (w/v) polyethylene glycol 6000 (PEG 6000, Merck) and incubation overnight at 4 °C. Precipitated T4Ps were pelleted by centrifugation at 12,000 rcf for 30 minutes at 4 °C. The resulting pellet was resuspended in buffer B and dialysed overnight against buffer B at 4 °C to remove PEG 6000. The presence of T4Ps was verified by SDS-PAGE followed by Coomassie visualisation of proteins and mass spectrometry (MRC-LMB Mass Spectrometry Facility, see Figure S1).

### Cryo-EM and cryo-ET sample preparation

For cryo-EM grid preparation of purified T4Ps, 2.5 µl of the sample was applied to a freshly glow-discharged Quantifoil R 3.5/1 Cu/Rh 200 mesh grid and plunge-frozen into liquid ethane using a Vitrobot Mark IV (ThermoFisher) at 100% humidity at an ambient temperature of 10 °C. For tomography of PAO1 Δ*pilT* cells, a bacterial lawn from an overnight LB agar plate incubated at 37 °C was resuspended in phosphate buffered saline (PBS), and 10 nm gold fiducials (CMC Utrecht) were added prior to plunge-freezing.

### Cryo-EM and cryo-ET data collection

Single particle cryo-EM data was collected on a Titan Krios G2 microscope (ThermoFisher) operating at an acceleration voltage of 300 kV, equipped with a Falcon 4i direct electron detector (ThermoFisher). Images were collected using a physical pixel size of 0.824 Å. For helical reconstruction of the T4P, movies were collected as 40 frames, with a total dose of 40 electrons/Å^2^, using a range of nominal defoci between -1 and -2.5 µm. For the T4P single-particle dataset, 5,679 movies were collected. Tomography data of PAO1 Δ*pilT P. aeruginosa* cells was collected on a Titan Krios G3 microscope (ThermoFisher) operating at an acceleration voltage of 300 kV, fitted with a Quantum energy filter (slit width 20 eV) and a K3 direct electron detector (Gatan). Cryo-ET tilt series were collected using a grouped dose-symmetric tilt scheme as implemented in SerialEM^62^, with a total dose of 175.5 electrons/Å^2^ per tilt series, nominal defocus of -8 µm, and with ±60° tilts of the specimen stage at 1° tilt increments. Tilt series images were collected using a physical pixel size 3.42 Å.

### Cryo-EM processing

Helical reconstruction of the T4P was performed in RELION 4.0^49,63–65^ (Fig. S2). Movies were motion-corrected using the RELION 4.0 implementation of MotionCor2^66^, and CTF parameters were estimated using CTFFIND4^67^. Initial helical symmetry of T4P used was obtained from previous reports in literature^24^. Three-dimensional (3D) classification was used to identify a subset of particles that supported refinement to 3.17 Å resolution. For final refinement, CTF multiplication was used for the final polished set of particles^47–49^. Symmetry searches were used during reconstruction, resulting in a final rise of 10.17 Å and a right-handed twist per subunit of 87.39°. Resolution was estimated using the gold-standard Fourier Shell Correlation (FSC) method as implemented in RELION 4.0. Local resolution measurements (Figure S3) were also performed using RELION 4.0.

### Model building and refinement

Manual model building of the PilA subunit was performed in Coot^68^. An AlphaFold2^69^ model was fit into the cryo-EM density as a rigid body. Due to the more flexible linker region, the α-helical parts and β-sheet were initially treated separately. Residues of the homology model that were inconsistent with the density were deleted and manually rebuilt. The initial model was subjected to real-space refinement against the cryo-EM map within the Phenix package^70,71^. Twenty-three subunits of PilA were built and used for final refinement. Non-crystallographic symmetry between individual PilA subunits was applied for all refinement runs. Model validation including map-vs-model resolution estimation was performed in Phenix (version 1.20).

### Tomogram reconstruction

Tilt series alignment via tracking of gold fiducials was performed using the IMOD^72^ implementation in RELION 5. Tomograms were reconstructed in RELION 5 and denoised using CryoCARE^73^ for visualisation purposes.

### Data visualisation and quantification

Atomic structures were visualised in ChimeraX^45^. Tomograms were visualised in IMOD^72^. Conservation analysis (Fig. 3h) was performed using the ConSurf server^74–76^. Hydrophobicity plots (Fig. S5) were calculated in ChimeraX using the in-built *mlp* function.

## Acknowledgments

This work was supported by the Medical Research Council, as part of United Kingdom Research and Innovation (also known as UK Research and Innovation) [Programme MC_UP_1201/31 to T.A.M.B.]. T.A.M.B. would like to thank EPSRC (Grant EP/V026623/1), the European Molecular Biology Organization, the Wellcome Trust (Grant 225317/Z/22/Z), the Leverhulme Trust, and the Lister Institute for Preventative Medicine for support. H.O. and I.C. were supported by EMBO Postdoctoral Fellowships (ALTF 1076-2023 to H.O. and ALTF 92-2022 to I.C.). The authors would like to thank Prof. Urs Jenal and Dr. Benoit-Joseph Levantie for sharing the *pilT* deletion strain, and Andriko von Kügelgen for help with the cryo-EM data collection. We acknowledge the MRC-LMB electron microscopy facility for help with sample preparation and data collection and the MRC-LMB mass spectrometry facility for assistance with mass spectrometry analysis of the sample.

## Author Contributions

H.O. and T.A.M.B. designed research. H.O., J.B., Z.W., A.K.T. and T.A.M.B. performed research. H.O., J.B., Z.W., A.K.T., I.C. and T.A.M.B. analysed data. H.O., J.B., Z.W., A.K.T. and T.A.M.B. wrote the manuscript.

## Supplementary Figures

**Figure S1:**
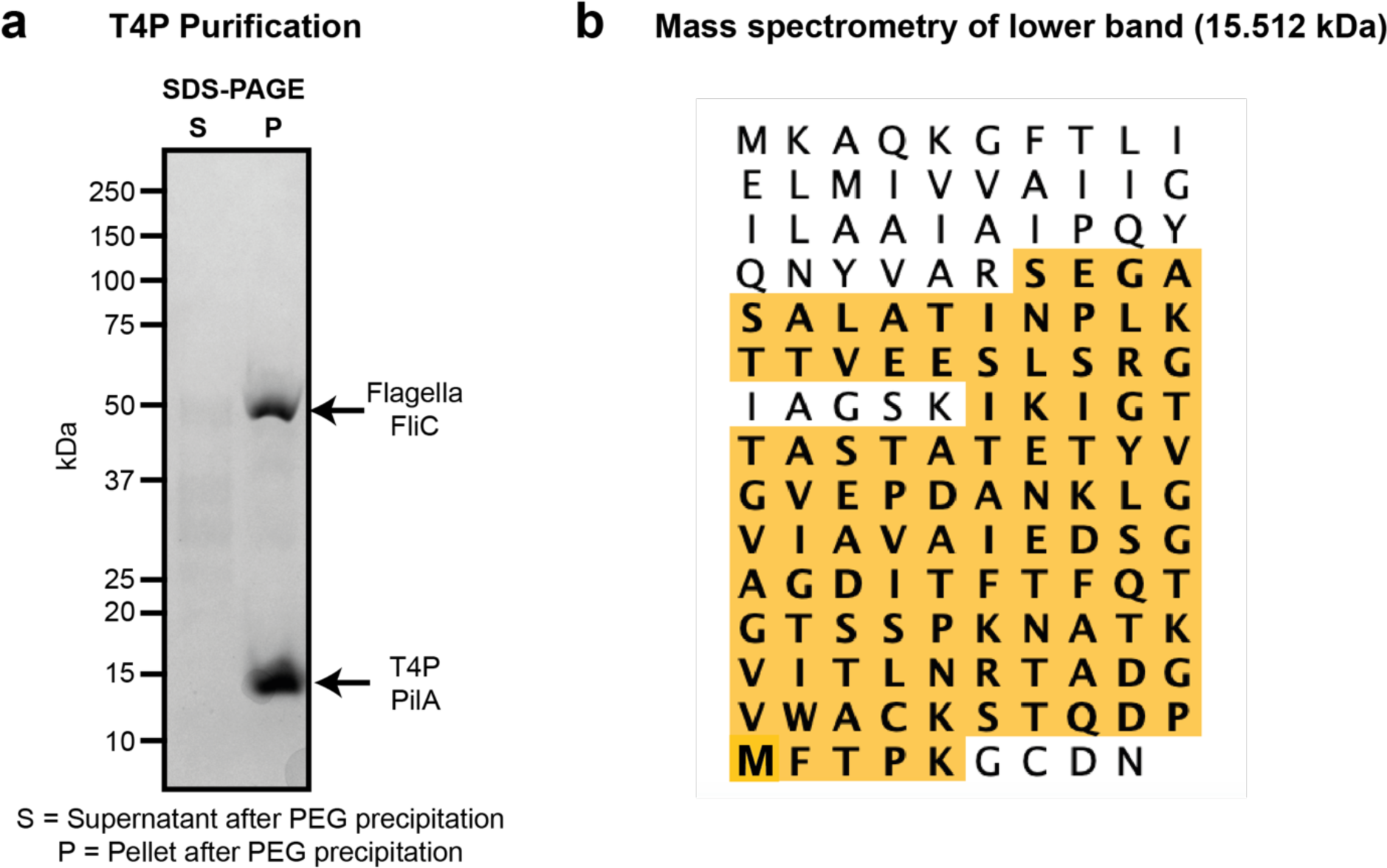
Purification of *P. aeruginosa* PAO1 T4P. **a** SDS-PAGE analysis of the final specimen after PEG precipitation (see Methods). Two bands, corresponding to FliC (flagella) and T4P PilA, are present at the expected molecular weights. The positions of molecular weight markers are shown on the left. **b** Peptide-fingerprinting mass spectrometry result of the T4P PilA band. The detected peptides are highlighted in the sequence, confirming the presence of T4P PilA in our sample.

**Figure S2:**
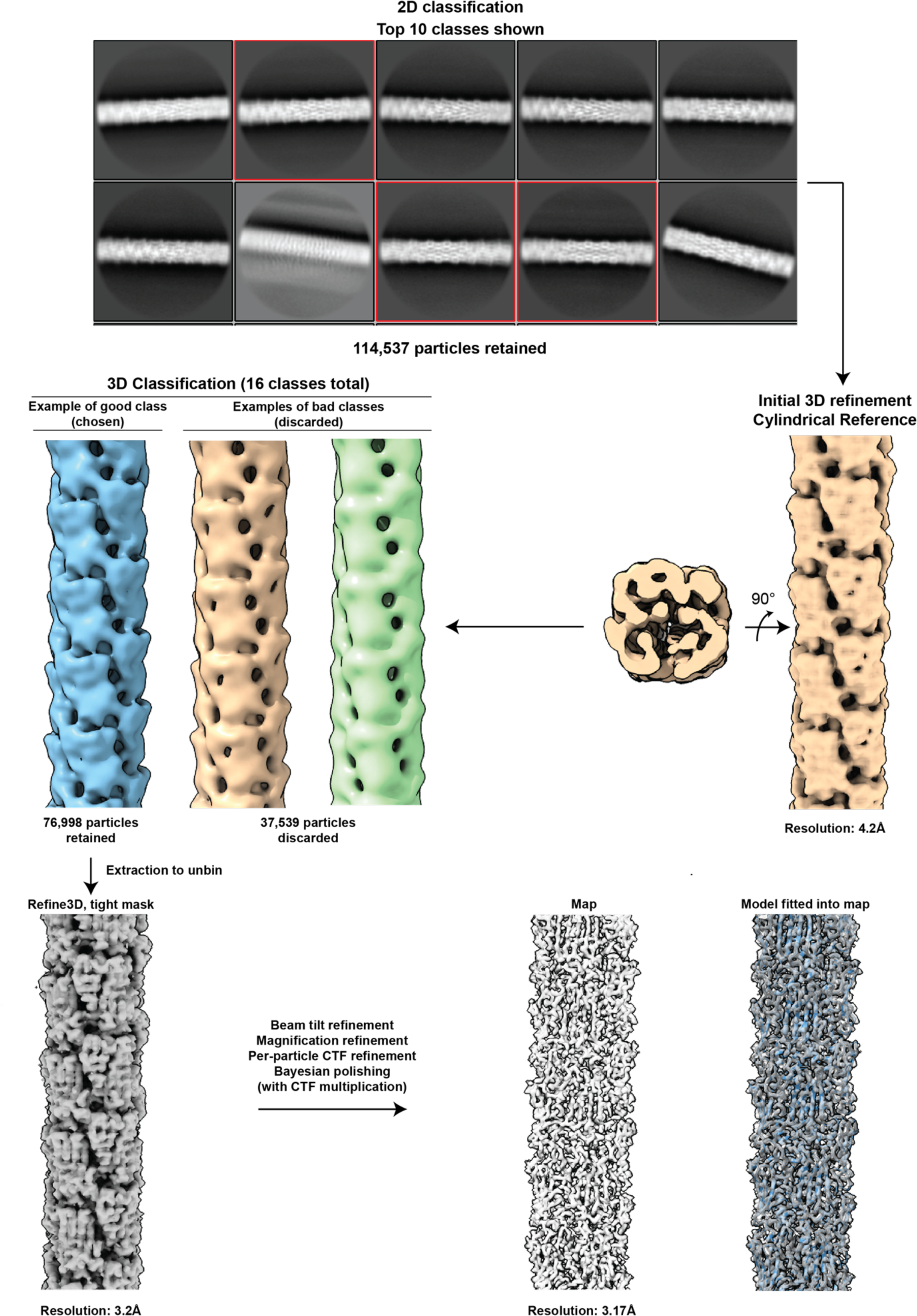
Cryo-EM image processing workflow in RELION 4.0. After particle picking and 2D classification (top 10 populated classes shown), the three best classes (114,537 particles, marked in red) were selected and used for initial 3D refinement yielding a 4.2 Å-resolution pilus structure. Following that, 3D classification was performed resulting in a final set of 76,998 particles, which were used for further 3D refinement. Further refinements and polishing were performed, resulting in the final 3.17 Å-resolution map.

**Figure S3:**
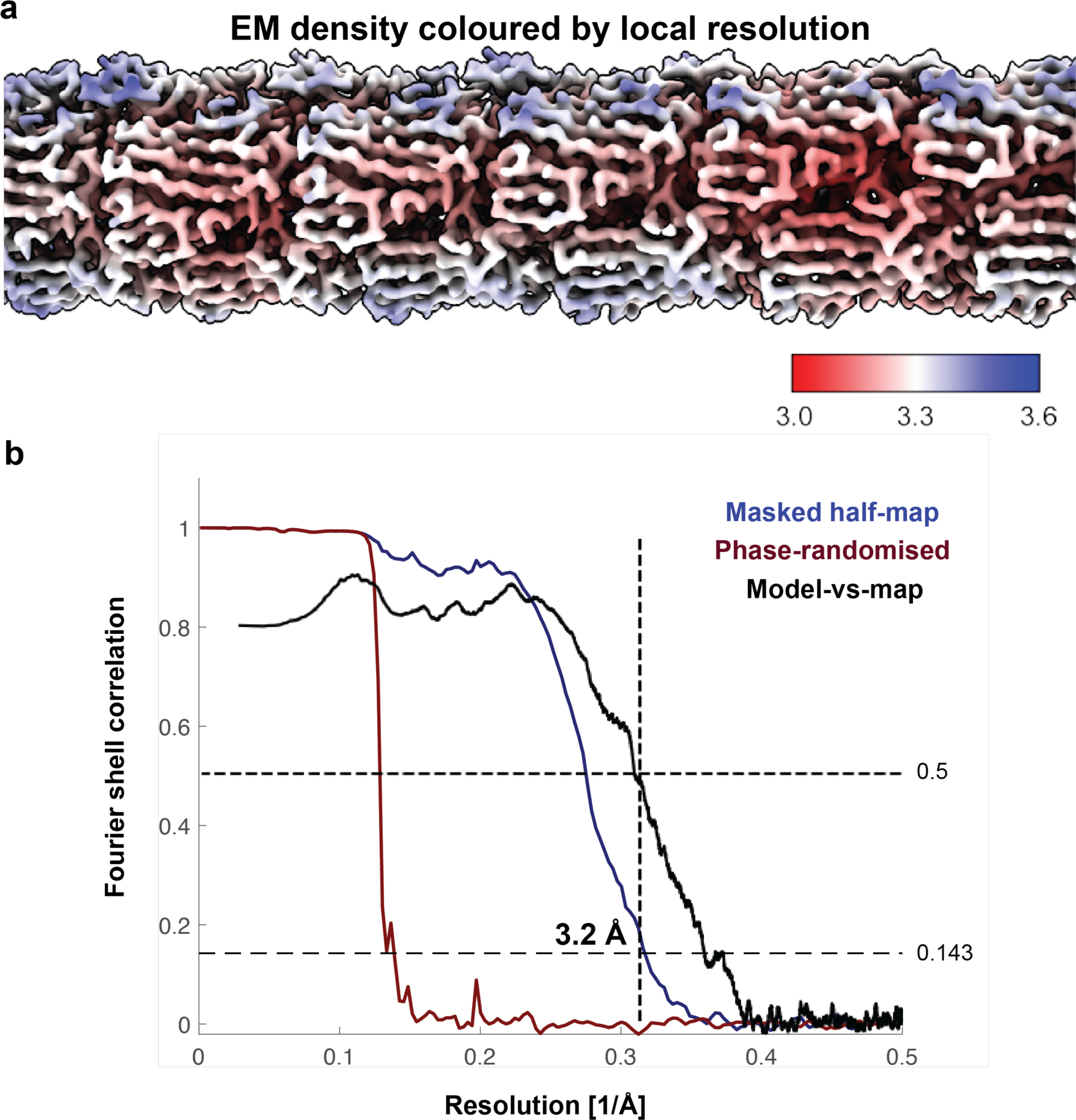
Resolution estimation of the map. **a** Cryo-EM density map coloured by local resolution (calculated in RELION 4.0). The slightly lower local resolution on the outside of the T4P is indicative of a higher degree of structural flexibility of the β-sheet domain compared to the α-helical part of the pilin. **b** Solvent-corrected masked half-map FSC (blue, gold-standard), phase-randomised half-map FSC (red), and model-vs-map FSC (black).

**Figure S4:**
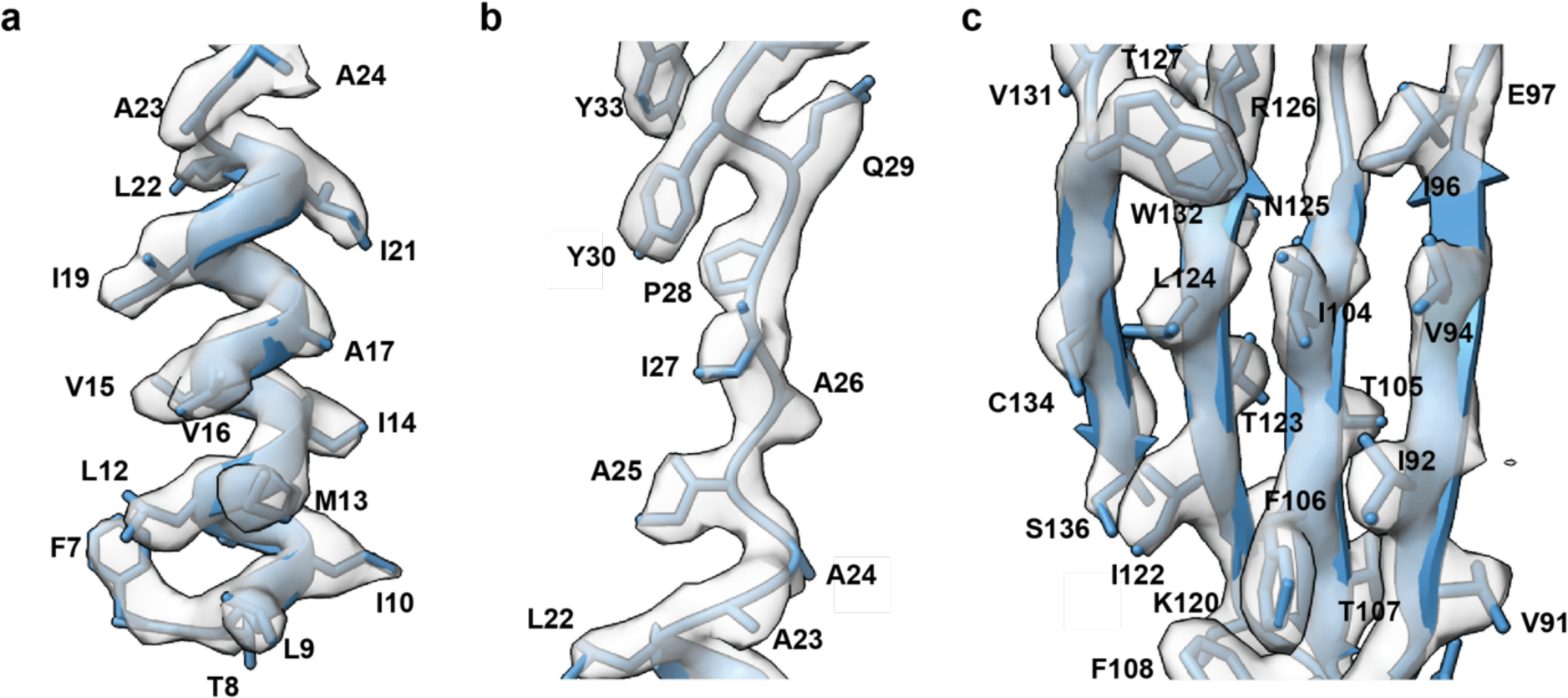
Side chain densities seen in the cryo-EM map. a-c. Models of the lower part of the N-terminal helix (residues 7-24, **a**), the ‘melted’ linker segment (residues 23-30, **b**) and the β-sheet (residues 91-136, **c**) fitted into the cryo-EM density. The side chains are clearly resolved in the map.

**Figure S5:**
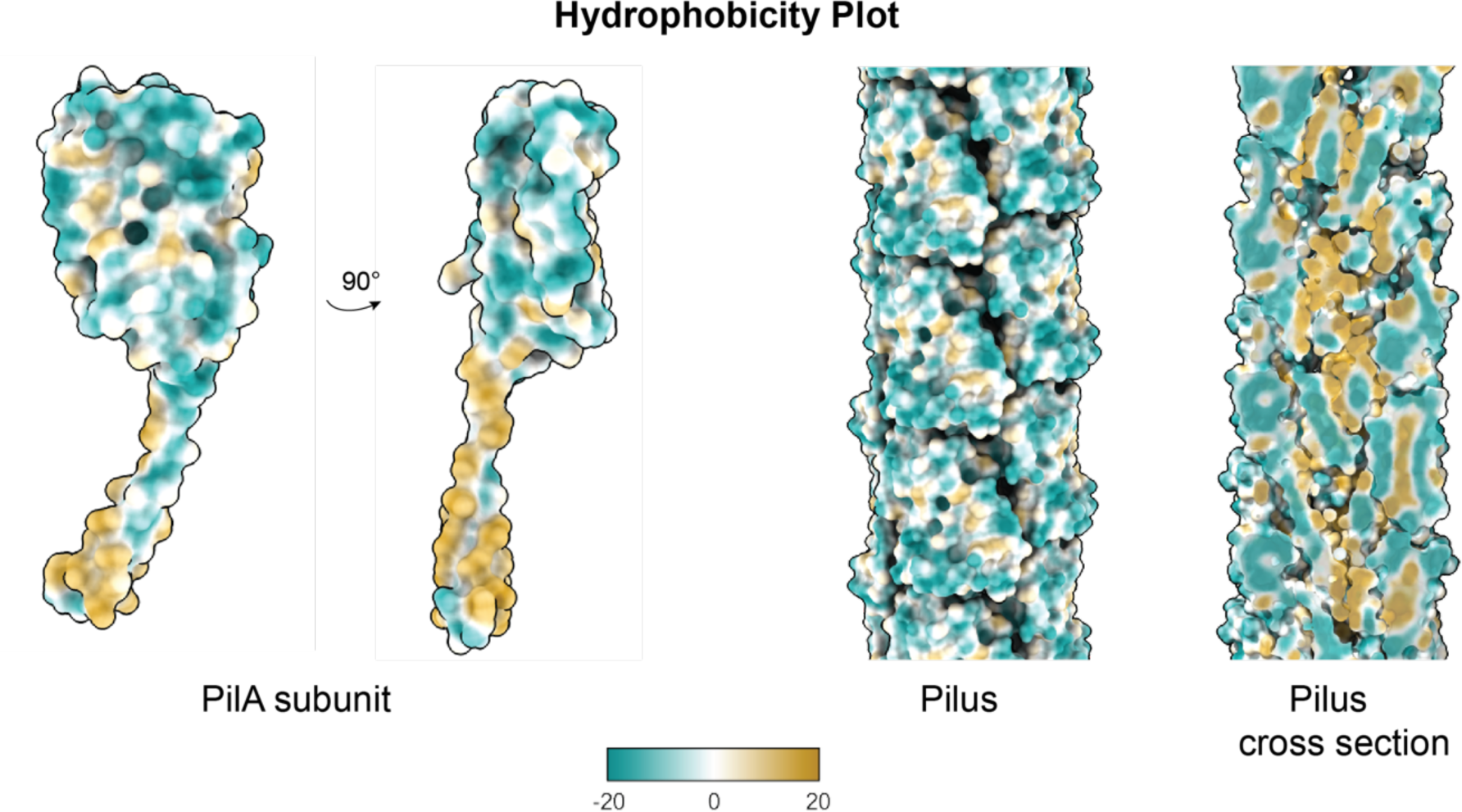
Hydrophobic properties of the *P. aeruginosa* PAO1 Type IV pilin and pilus. Hydrophobicity of the *P. aeruginosa* PilA subunit and the fully assembled T4P. The subunit is shown in two orientations related to each other by a 90° rotation (left). For the assembled pilus (right), both the outer surface and a cross section through the centre are shown. The models have been coloured by molecular lipophilicity potential using the *mlp* function in ChimeraX^45^.

**Figure S6:**
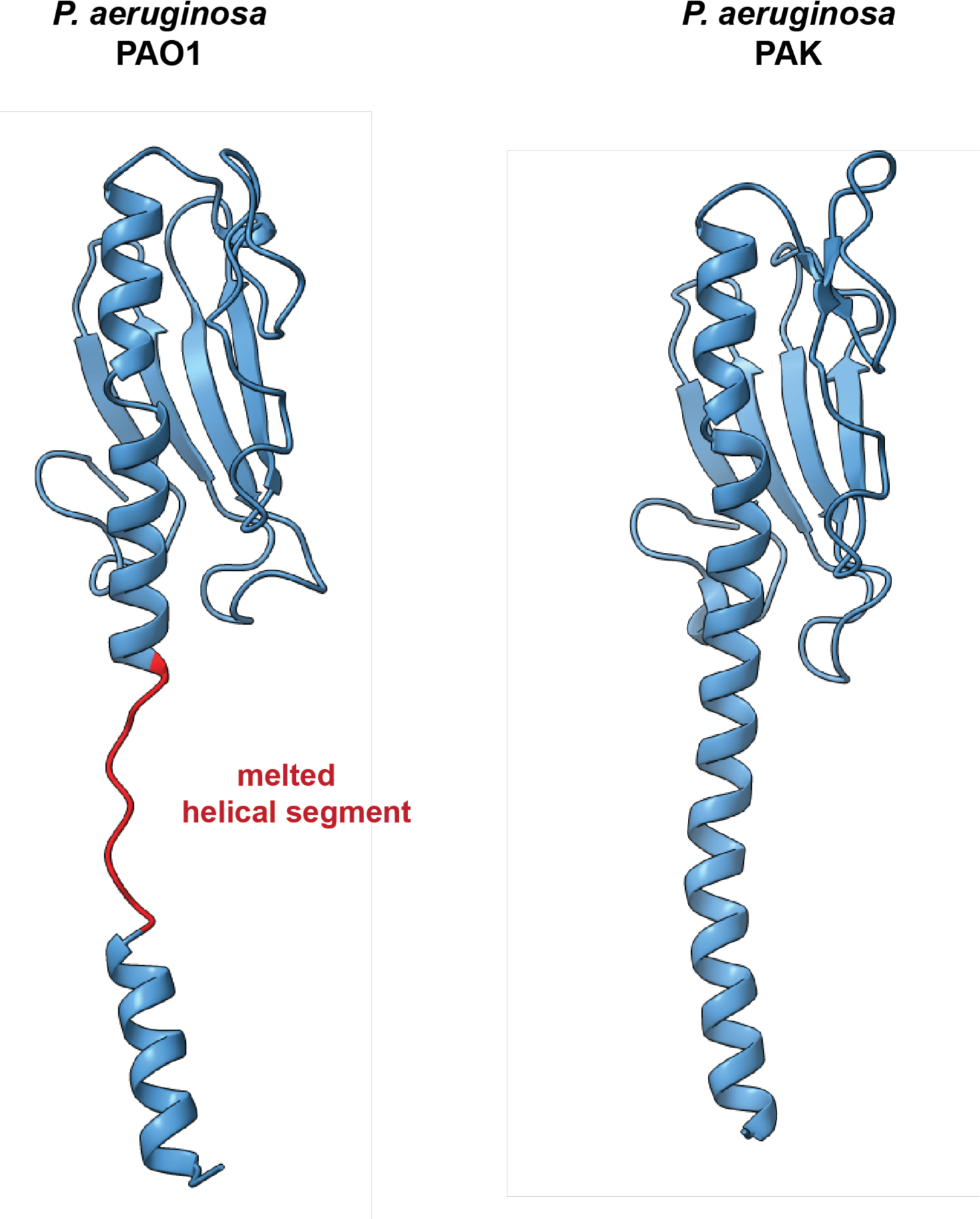
Comparison of the PAO1 PilA cryo-EM structure to the PAK PilA X-ray crystallography structure. Side-by-side comparison of the PAO1 PilA (this study, left) with the PAK PilA (PDB: 1OQW, right) demonstrating the absence of the melted helical segment in the latter.

## Supplementary Tables

**Table S1:** Cryo-EM data acquisition and processing statistics.

|  |  |
| --- | --- |
| **Data collection and processing** | PilA ΔpilT [EMDB 50025, PDB 9EWX] |
| Microscope | Krios Titan G2 |
| Magnification | 96,000 |
| Voltage (kV) | 300 |
| Electron exposure (e<sup>-</sup>/Å<sup>2</sup>) | 40 |
| Defocus range (μm) | -1 to -2.5 |
| Pixel size (Å) | 0.824 |
| Symmetry imposed | Helical, final: 87.39° 10.17 Å |
| Initial particle images (no.) | 114,537 |
| Final particle images (no.) | 76,998 |
| Map resolution (Å) | 3.17 |
| FSC threshold | 0.143 |
| Map resolution range (Å) | 3.0-3.7 |
| **Refinement** |  |
| Initial model used | AlphaFold2 |
| Model resolution (Å) | 2.6/2.8/3.2 |
| FSC threshold | 0/0.143/0.5 |
| Model resolution range (Å) | n/a |
| Map sharpening *B* factor (Å<sup>2</sup>) | -42.7 |
| Model composition |  |
| Non-hydrogen atoms | 23,943 |
| Protein residues | 3,289 |
| Ligands | 0 |
| *B* factors (Å<sup>2</sup>) |  |
| Protein | 28.74 |
| Ligand | n/a |
| R.m.s. deviations |  |
| Bond lengths (Å) | 0.003 |
| Bond angles (°) | 0.554 |
| Validation |  |
| MolProbity score | 1.22 |
| Clashscore | 2.8 |
| Poor rotamers (%) | 0 |
| Ramachandran plot |  |
| Favored (%) | 97.16 |
| Allowed (%) | 2.84 |
| Disallowed (%) | 0 |

**Table S2:** T4P from various organisms.

|  | *Pseudomonas aeruginosa* | *Neisseria gonorrhoeae* | *Escherichia coli* | *Geobacter sulfurreducens* | *Thermus thermophilus* | *Myxococcus xanthus* | *Pyrobaculum arsenaticum* | *Monoderm bacterium* |
| --- | --- | --- | --- | --- | --- | --- | --- | --- |
| Pilin sequence length | 143 | 156 | 140 | 104 | 125 | 208 | 140 | 137 |
| Pilus diameter | 51Å | 62Å | 60Å | 65Å | 70Å | 70Å | 80Å | 70Å |
| Helical rise | 10.17Å | 10.1Å | 11.2Å | 10.4Å | 9.33Å | 10.0Å | 5.3Å | 11Å |
| Helical twist | 87.39° | 100.8° | 96° | 89.1° | 92.5° | 100.7° | 101.7° | 93° |
| Helical pitch | 41Å | 36Å | 42Å | 42Å | 36Å | 36Å | 19Å | 43Å |
| PDB ID | 9EWX | 5VXX | 6GV9 | 6VK9 | 6XXD | 8TJ2 | 6W8U | 8PFB |

## Supplementary Movie Legends

**Movie S1: Cryo-EM structure of the *P. aeruginosa* PAO1 T4P.**

A 3.2 Å resolution cryo-EM density map of the T4P is shown, which was used to build an atomic model of the main pilus-forming subunit PilA (surface depiction and ribbon diagrams are shown).

